# A novel mesocosm set-up reveals strong methane emission reduction in submerged peat moss *Sphagnum cuspidatum* by tightly associated methanotrophs

**DOI:** 10.1101/536268

**Authors:** Martine A. R. Kox, Alfons J. P. Smolders, Daan R. Speth, Leon P. M. Lamers, Huub J. M. Op den Camp, Mike S. M. Jetten, Maartje A. H. J. van Kessel

## Abstract

Wetlands present the largest natural sources of methane (CH_4_) and their potential CH_4_ emissions greatly vary due to the activity of CH_4_-oxidizing bacteria associated with wetland plant species. In this study, the association of CH_4_-oxidizing bacteria with submerged *Sphagnum* peat mosses was studied, followed by the development of a novel mesocosm set-up. This set-up enabled the precise control of CH_4_ input and allowed for monitoring the dissolved CH_4_ in a *Sphagnum* moss layer while mimicking natural conditions. Two mesocosm set-ups were used in parallel: one containing a *Sphagnum* moss layer in peat water, and a control only containing peat water. Moss-associated CH_4_ oxidizers in the field could reduce net CH_4_ emission up to 93%, and in the mesocosm set-up up to 31%. Furthermore, CH_4_ oxidation was only associated with *Sphagnum*, and did not occur in peat water. Especially methanotrophs containing a soluble methane monooxygenase enzyme were significantly enriched during the 32 day mesocosm incubations. Together these findings showed the new mesocosm setup is very suited to study CH_4_ cycling in submerged *Sphagnum* moss community under controlled conditions. Furthermore, the tight associated between *Sphagnum* peat mosses and methanotrophs can significantly reduce CH_4_ emissions in submerged peatlands.

## Introduction

Methane (CH_4_) has a 25 times higher Global Warming Potential (GWP) than carbon dioxide (CO_2_; on a 100 year time scale) and is the second most important greenhouse gas (GHG), contributing for about 16% to global warming [1, 2]. CH_4_ in the atmosphere originates from both natural and anthropogenic sources. Wetlands are the largest natural CH_4_ source, emitting an estimated 167 Tg CH_4_ yr^−1^ into the atmosphere [3], indicating an imbalance between CH_4_ production and CH_4_ consumption by methanotrophs. Climate change has the potential to further stimulate the emission of CH_4_ from (especially artic) wetlands [4]. Therefore, it is important to understand sources, sinks and microbial transformations of CH_4_ in wetland ecosystems.

CH_4_ cycling in peat ecosystems is affected by peat degradation and subsequent restoration [5–7]. Restored (rewetted) sites appear to emit more CH_4_, indicating that restored conditions stimulate methanogenesis, and that methanotrophy cannot keep up. One well-known factor controlling CH_4_ cycling in wetlands is the water-table [8, 9]. The CH_4_ emission from rewetted peatlands remains low when the water table remains well below the field surface. However, when the water-table rises, CH_4_ emission strongly increases [10, 11]. As an example, the Mariapeel peatland in The Netherlands has been drained for many years, leading to severe drought. The peatland was rewetted again for restoration purposes, which resulted in a strong decrease of CO_2_ emissions that originated from the aerobic oxidation of organic material, whereas the emission of the much stronger greenhouse gas CH_4_ emission strongly increased [10]. The CH_4_ emission in rewetted peatlands seems to be strongly reduced by development of (aquatic) *Sphagnum* mosses, which harbor CH_4_-oxidizing microorganisms [6, 10, 12]. It is, however, challenging to study CH_4_ dynamics in primary stages of peat development (either restored/natural) without disturbing the site. Furthermore, also abiotic factors such as temperature, water quality and light availability on site cannot be controlled as well as in the laboratory, making experimental work and predictions about peat development and CH_4_ cycling at least cumbersome.

As mentioned above, CH_4_ emissions are caused by an imbalance between CH_4_ production and consumption. The CH_4_ emitted by peatlands is mainly produced by methanogenic Archaea [13]. In the anaerobic, submerged peat layers that are devoid of electron acceptors other than CO_2_, methanogens produce CH_4_ from a limited number of substrates and/or in syntrophic interaction with other anaerobes that degrade organic carbon (C). However, not all of the CH_4_ produced reaches the atmosphere, due to methanotrophs that oxidize CH_4_ to CO_2_ [14, 15]. The oxidation of CH_4_ is performed both aerobically (e-acceptor: O_2_) by CH_4_-oxidizing bacteria (MOB), and anaerobically (AOM) by Archaea and bacteria (e-acceptors: nitrite, nitrate, metal-oxides, humic acids, and sulfate [16]). Both aerobic and anaerobic CH_4_ oxidation contribute to the reduction of CH_4_ emissions from peatlands [12, 17–19]. Within the MOB the enzyme methane monooxygenase (MMO) is responsible for the oxidation of CH_4_ to methanol. The majority of MOB have a copper containing, membrane bound form of MMO (pMMO) [20]. In addition, a small fraction of the MOB also has a soluble form of MMO (iron containing sMMO) [20]. The sMMO seems to be only expressed when copper limitation is experienced and has a less restricted substrate specificity than pMMO [20]. Peatland methanotrophs typically possess both pMMO and sMMO [12, 21–23], which can be targeted via the *pmoA* and *mmoX* genes encoding one of the subunits, respectively. Some peatland and marine methanotrophs are unique in that they only possess sMMO [23–27]. Also *mmoX* transcripts indicate that sMMO is an active enzyme in peatlands [28], although its importance is not yet well understood.

Studies have shown that aerobic CH_4_ oxidation is most prominent in submerged *Sphagnum* mosses in a range of peatlands [12, 29, 30]. Furthermore, the association between methanotrophs and *Sphagnum* was shown to be mutually beneficial Raghoebarsing et al. [31]. The methanotrophs convert CH_4_ into CO_2_, thereby relieving part of the CO_2_ limitation that *Sphagnum* mosses experience [32] especially under submerged conditions [12, 31]. The aerobic MOB in return benefit from O_2_ produced and shelter provided by the moss [12].

Molecular surveys showed that several CH_4_-oxidizing bacteria are present in *Sphagnum* dominated peatlands. *Alphaproteobacterial* methanotrophs typically dominate in 16S rRNA gene libraries over the other methanotroph-containing (sub)phyla *Gammaproteobacteria* and *Verrucomicrobia (Methylacidiphilaceae* [33–35]. Within the *Alphaproteobacteria* especially methanotrophs of the family *Methylocystaceae (Methylocystis spp.)* and the acidophilic methanotrophs of the family *Beijerinckiaceae (Methylocella, Methyloferula, Methylocapsa)* are often found and several of these have been isolated from peatlands [24–26, 36, 37]. Using Fluorescence *in situ* Hybridization (FISH) combined with confocal microscopy, *Alphaproteobacteria* have shown to be localized inside *Sphagnum* mosses, in the dead hyaline cells [38]. Furthermore, *Verrucomicrobia* including the class containing CH_4_ oxidizers, *Methylacidiphilae*, can make up 10% of the total microbial community associated with *Sphagnum*. However, the *Methylacidiphilae* found with *Sphagnum* mosses have not yet been coupled to CH_4_-oxidizing activity [34, 39, 40]. Their role in peatland C cycling has yet to be confirmed [23, 41–43].

The goal of this study was to design and test a new mesocosm set-up where a submerged *Sphagnum* community could be mimicked under fully controlled conditions. In this way, the irregularity and variability often encountered in field studies could be excluded. The new set-up was used to study the association between CH_4_ oxidizers and a layer of submerged *Sphagnum* mosses. We hypothesized that the submerged *Sphagnum* moss layer acts as a biofilter for CH_4_, thereby reducing CH_4_ emission to the atmosphere. Furthermore, it is expected that the CH_4_-oxidizing microorganisms are associated with *Sphagnum*, rather than the peat water. Monitoring of the CH_4_ flux throughout the mesocosm incubation, as well as CH_4_ batch assays and molecular analysis of 16S rRNA gene amplicons and qPCR on 16S rRNA, *pmoA* and *mmoX* showed that during the 32 days of incubation aerobic methanotrophs were highly active and enriched in the mesocosm.

## Materials & Methods

### Sampling site and field measurements

The sampling site was located in the Mariapeel (51°24’28.4”N, 5°55’8”E), a peat bog nature conservation area in the south of the Netherlands. This site was visited for measurements and sampling on 09/08/2017. Net diffusive gas fluxes of CO_2_ and CH_4_ were measured in the field using a fast greenhouse gas analyzer with cavity ringdown spectroscopy (GGA-24EP; Los Gatos Research, USA) connected to a Perspex chamber (15 cm in diameter). The chamber was put on top of the moss layer for 10 min to measure fluxes of CO_2_ and CH_4_. In total 3 independent measurements were taken within 2 m distance from each other. After removal of the peat moss layer measurements were repeated, after an equilibration period of 15 min. Submerged *Sphagnum cuspidatum* moss and water were collected after the measurements.

Upon arrival in the laboratory, 1 set of mosses was used to determine field activity, and another part was washed using sterile demineralized H_2_O. One fraction of water was used to determine field activity, the other fraction was filtered (2 − 5 nm, HF80S dialysis filter, Fresenius Medical Care, Homburg, Germany). All samples were stored at 4 °C (1 week) until the start of the incubation.

### Mesocosm design

The mesocosm consists of a glass cylinder with a diameter of 12 cm and a height of 54 cm, to which a separate reservoir is connected (see Supplementary Figs. 1 and S1). The total reservoir volume is 0.5 L, the connector tube volume is 0.07 L and the total column volume is 6.11 L. The water level in the mesocosms was maintained at 5.09 L, leaving a headspace of 1.02 L in the column. The column headspace was closed throughout the day using a greased lid with sampling port. Several sampling ports (in the reservoir, cylinder headspace and in the cylinder at 10, 20, 30, 35 and 40 cm height) allow for sampling of either the gas or water phase. Throughout the mesocosm incubation all sampling ports were closed off using boiled, red butyl rubber stoppers and capped using metal crimp caps.

**Fig. 1.**
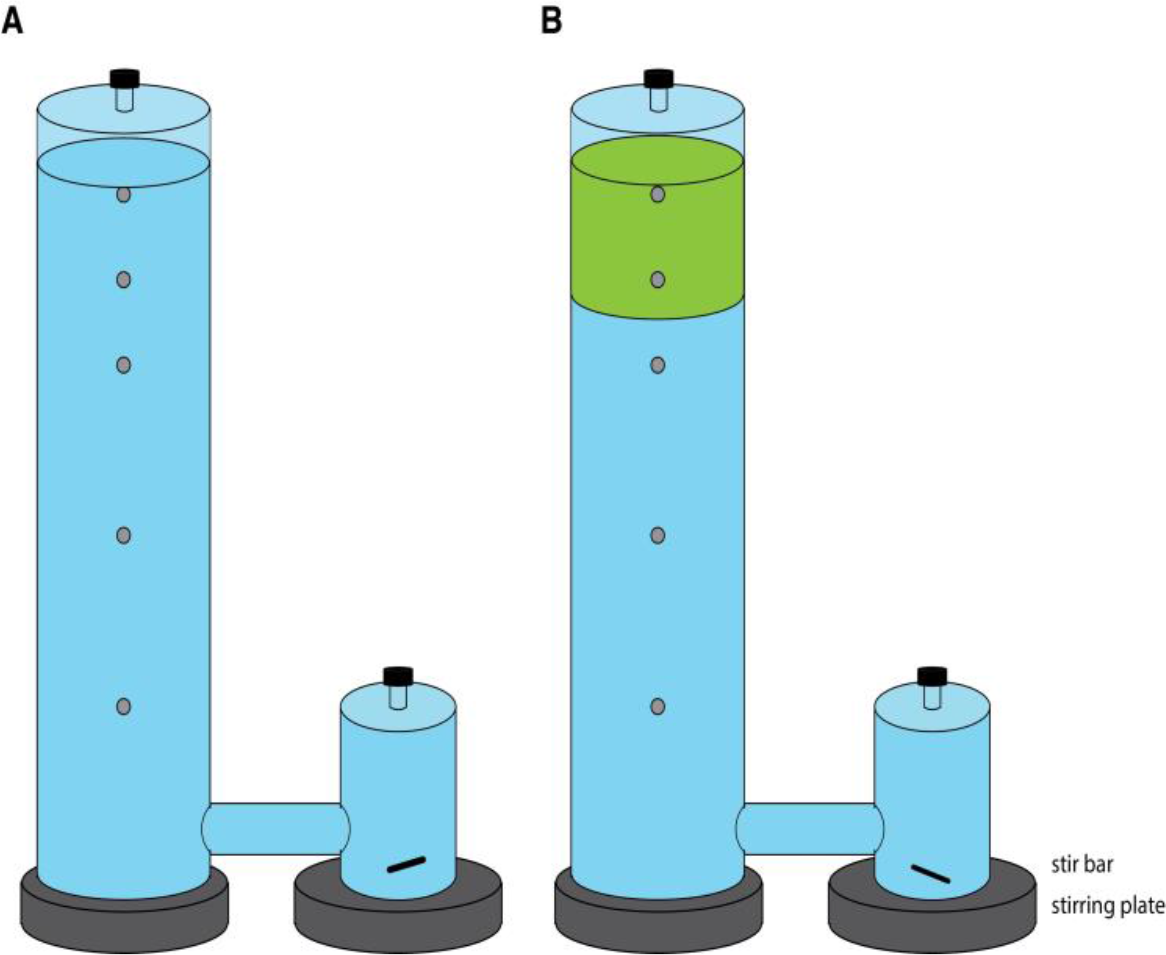
Schematic set up of mesocosm incubation with in **A.** control mesocosms containing only filtered peat water (blue) and in **B.** the moss mesocosms, containing sphagnum moss layer (green) in filtered peat water.

### Mesocosm incubation

The mesocosms were autoclaved prior to use. Two mesocosms were simultaneously incubated for this experiment. A moss mesocosm, containing 100 *Sphagnum cuspidatum* plants (6 cm length, 120 g fresh weight) in filtered peat water (5.09 L), and a control mesocosm which contained only filtered peat water (5.09 L). Both mesocosms had an acclimatization period of 7 days prior to sampling.

The CH_4_ was added via the reservoir headspace and dissolved into the water by stirring with a 2 cm magnetic stir bar at 250 rpm. Throughout the week, lids were opened each morning for 1 h to allow aeration, after which they were closed for the rest of the day. The CH_4_ supply in the reservoir headspace was replaced daily, directly after aeration, with a mixture of 50ml 99% CH_4_ and 5 ml CO_2_. The mesocosm experiment was performed twice, each time for 32 days. Incubations were performed at room temperature. The light regime consisted of 16 h daylight (150 μmol m^−2^ s^−1^ photosynthetically active radiation at vegetation level) and 8 h of darkness. Light was supplied on top of the mesocosm column, via 120 deep red/white LED lamps (Philips, Green-Power LED, Poland).

### Mesocosm CH_4_ fluxes

After the acclimatization period the fate of CH_4_ was followed through the mesocosm over time (0 - 32 days). To determine the concentration of CH_4_ in the headspace or the concentration of dissolved CH_4_ in water, gas and water samples were collected via the different sampling ports. A volume of 0.5 ml gas or 0.5 ml water was taken and injected into a closed 5.9 ml Exetainer vial (Labco, Lampeter, UK). The concentration of CH_4_ in the headspaces of the reservoir and the column were determined by taking samples directly after closing the column in the morning (0 h) and before opening the column for aeration again (23 h). The concentration of dissolved CH_4_ throughout the column was determined once a week, by sampling water at 4 different time points during the day (0 h, 3 h, 7 h, 23 h after closing the headspace).

The CH_4_ concentration in the Exetainers was measured at least 4 h after sampling to allow for equilibration between Exetainer headspace and liquid. The CH_4_ concentration was measured using a gas chromatograph with a flame-ionized detector and a Porapak Q column as described by De Jong et al. [4].

Net CH_4_ flux in the mesocosm was calculated as the change in CH_4_ concentration in the headspace of the mesocosm column for each day and divided by the surface area (0.01131 m^2^) of the mesocosm column.

### Potential CH_4_ oxidation rates

The CH_4_ oxidation rates were determined in triplicate in batch incubations prior to and after mesocosm incubation. Prior to the mesocosm incubation, both unwashed and washed moss (3 g fresh weight) as well as unfiltered and filtered porewater (12 ml) were placed into autoclaved 120 ml serum vials and closed with boiled, red-butyl rubber stoppers and metal crimp-caps. Each batch flask received 2 ml 99% CH_4_. The CH_4_ concentration in the headspace was followed in time as described for mesocosm CH_4_ fluxes.

At the end of the mesocosm experiment, potential CH_4_ oxidation rates were determined for the mosses from moss mesocosm and for porewater from both the moss and control mesocosm. Samples were incubated as described above. Two sets of each 3 replicates were incubated, where one set was used to determine CH_4_ oxidation rates and the other set received the acetylene (6 ml 99.9% (C_2_H_2_)), an inhibitor of the CH_4_ monooxygenase enzyme, which was added after 10 h of incubation.

The concentrations of CH_4_ were calculated using a calibration curve that was measured daily. Ultimately, the CH_4_ concentrations were plotted over time, from which CH_4_ oxidation rates were calculated from the slope of the linear part of the graph.

### Elemental analysis water

Both unfiltered and filtered peat water was sampled and analyzed. The pH was measured and elemental composition was determined using the auto analyzer and the ICP-OES as explained before [34]. Dissolved CH_4_ in field porewater was determined by injection of 1 ml porewater into a closed Exetainer (5.9 ml), after 6 hours the headspace CH_4_ concentration was measured as described above. Data are shown in Supplementary Table S9.

### DNA extraction

DNA extraction was performed by grinding 5 g of mosses (fresh weight) using pestle and mortar and liquid nitrogen, after which DNA was extracted using the DNeasy Powersoil DNA extraction kit following manufacturers protocol (Qiagen Benelux B.V., Venlo, Netherlands). DNA quality was checked by gel electrophoresis (1% agarose gel in TBE buffer) and fluorometrically using the Qbit dsDNA HS Assay Kit (Invitrogen, Thermo Fisher, Carlsbad, CA).

### Amplicon sequencing and analysis

Barcoded Amplicon sequencing of the amplified V3-V4 region of the bacterial 16s rRNA gene (primers Bact-341f and Bact 785r [44]) was done using Illumina Miseq, performed by BaseClear B.V. (Leiden, the Netherlands). A total of 326 045 reads was obtained. The reads were quality filtered and analyzed using Mothur (v1.36.1), following the Illumina Standard Operating Procedure (SOP, accessed on May 8^th^ 2018, Kozich et al. 2013). Merged reads shorter than 400 bp were discarded, chimeras were removed using the UCHIME algorithm [46] and the remaining sequences were clustered at 97% identity. The resulting OTUs were classified based on the SILVA v132 16s rRNA gene non-redundant database (SSURef_99_v132_SILVA). Next, non-target sequences (Chloroplasts, Mitochondria, unknown, Archaea and Eukaryota) were removed from the dataset. See Supplementary Tables S1 and S2 for full overview of read processing. The output was analyzed with R (version 3.4.0 by the R Development Core Team [47]) and Rstudio v1.1.456 [48] using the packages Phyloseq [49] and vegan [50]. Singletons were removed, and read libraries of all samples were rarefied by random subsampling (seed: 12345) to 6 500 reads per sample (Rarefaction curves are depicted in Supplementary Fig. S2). A PcoA plot (Supplementary Fig. S4) was created using Phyloseq, and based upon Bray-Curtis dissimilarity matrix on rarefied data. All sequencing data can be accessed in GenBank NCBI BioProject PRJNA517391.

### Quantitative PCR

Copy numbers of the Bacterial 16S rRNA gene (for all primers see Table S3; Bact 341f - Bact 785r; Klindworth et al. 2013), as well as functional genes *pmoA* (primers A189f-A682r; Holmes et al. 1995) and *mmoX* (mmoX1-mmoX2; Miguez et al. 1997) were quantified using a qPCR approach. The qPCR reaction mix consisted of PerfeCTA Quanta master mix (Quanta Bio, Beverly, MA) and 0.5 ng sample DNA and 1μl of each primer (10 μM). In negative controls DNA was replaced by sterilized milli-Q water. The qPCR reaction mix was loaded in triplicate into a 96-well optical PCR plates (Bio-Rad Laboratories B.V., Veenendaal, The Netherlands), closed with an optical adhesive cover (Applied Biosystems, Foster City, CA) and reactions were performed with a C1000 Touch thermal cycler equipped with a CFX96 Touch™ Real-Time PCR Detection System (Bio-Rad Laboratories B.V., Veenendaal, The Netherlands). Standard curves were obtained via 10-fold dilution series of a PGEM T-easy plasmid (Promega, Madison, WI) containing the target gene. The data was analyzed using Bio-Rad CFX Manager version 3.0 (Bio-Rad Laboratories B.V., Veenendaal, The Netherlands). Triplicate analysis per samples were averaged prior to statistical analysis.

### Statistics

The CH_4_ flux in the field and in the mesocosm, CH_4_ oxidation rates in batch and qPCR data were analyzed using R version 3.4.0 by the R Development Core Team [47]. In order to allow for parametrical statistical tests, Shapiro-Wilk’s test was used on the residual (stats-package) to test the normality of the data and Levene’s test (car-package) was used to test for homogeneity of variance. If assumptions of tests were not met, data was log-transformed (ln), which was the case for the field CH_4_ flux data. A paired T-test was used to test whether the net CH_4_ flux in the field was affected by the presence of moss (moss field / moss removed). Differences between material (moss/peatwater) in the potential CH_4_ oxidation activity prior to mesocosm incubation was tested using a non-parametric Kruskal Wallis tests. Within each material (moss/peatwater) the effect of treatment (field / washing or filtering) was tested using an independent T-test.

Differences between mesocosms (moss / control), material (moss / peat water) and inhibitor (yes/no) in the potential CH_4_ oxidation activity after mesocosm incubation, were tested using a 3-way Anova, followed by a Tukey HSD post-hoc test. Differences in copy number between each moss sample (Moss Field/Moss Washed/Moss incubated) within each target gene (*16S rRNA/mmoX*/*pmoA*) was analyzed using a one-way Anova, followed by a Tukey HSD post-hoc test. Here, the data for *16S rRNA* gene and *mmoX* gene were log-transformed (ln) prior to analysis.

## Results

### Field CH_4_ flux

To estimate diffusive CH_4_ emissions in the field, flux chamber measurements were carried out in plots with submerged *Sphagnum* mosses before and after removal of the moss layer. The CH_4_ emission in the field situation with the submerged *Sphagnum* moss layer resulted in a net total of 4.1 ± 2.1 mmol CH_4_ m^−2^ day^−1^ (mean ± SEM, n=3; Fig. 2). Removal of the *Sphagnum* moss layer significantly increased the net CH_4_ emission (t_(2)_ = −6.1, p < 0.05) to a total of 60 ± 32 mmol CH_4_ m^−2^ day^−1^ (Fig. 2).

**Fig. 2.**
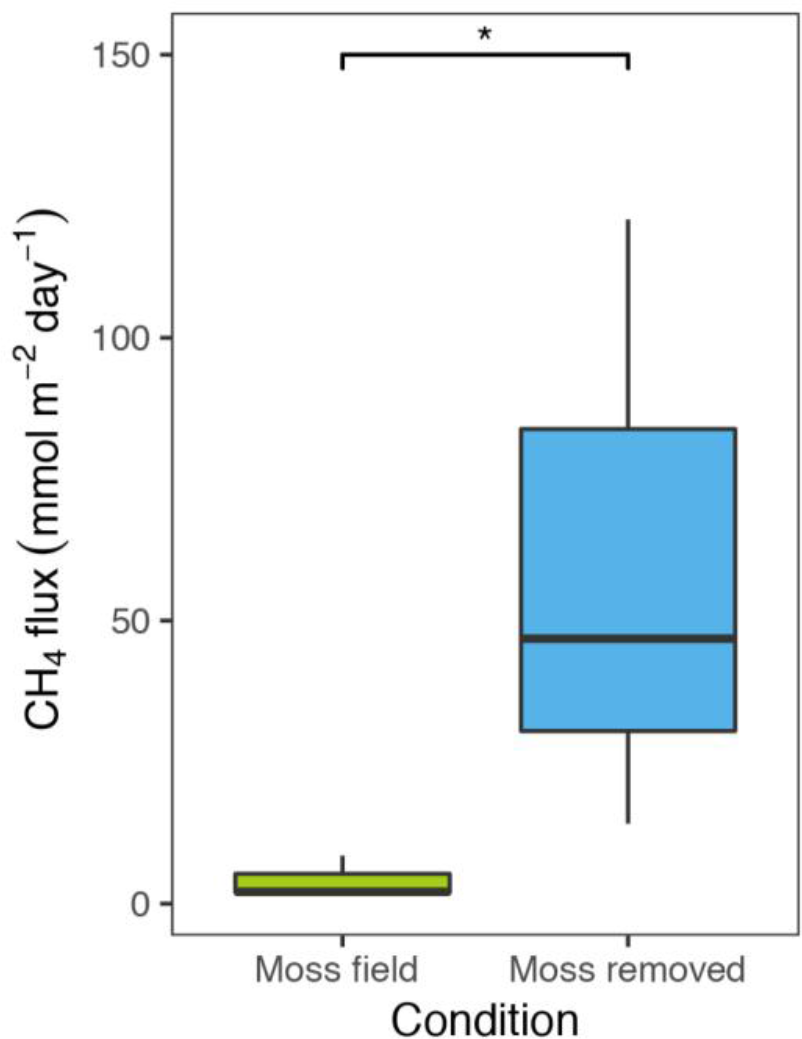
Net CH_4_ flux (mmol CH_4_ m^−2^ day^−1^) measured in the field with *Sphagnum* moss layer present (green, n=3) and after moss removal (blue, n=3). Error bars indicate the standard error of the mean.

### Methane oxidation activity prior to mesocosm incubation

The CH_4_ oxidation rates associated with the *Sphagnum* moss and peat water were determined prior to the incubation in the mesocosm, using batch assays (Fig. 3). *Sphagnum* mosses showed much higher CH_4_ oxidation rates (average rate mosses 143 ± 17 μmol g DW^−1^ day^−1^, Fig. 3) compared to peat water, which had virtually no activity (0.05 ± 0.06 μmol g DW^−1^ day^−1^; χ^2^ = 7.5, p < 0.01, Supplementary Fig. S5). Washing of the *Sphagnum* mosses was reduced the CH_4_ oxidation rate 121 μmol g DW^−1^ day^−1^; t_(2)_ = 1.5, p > 0.05).

**Fig. 3.**
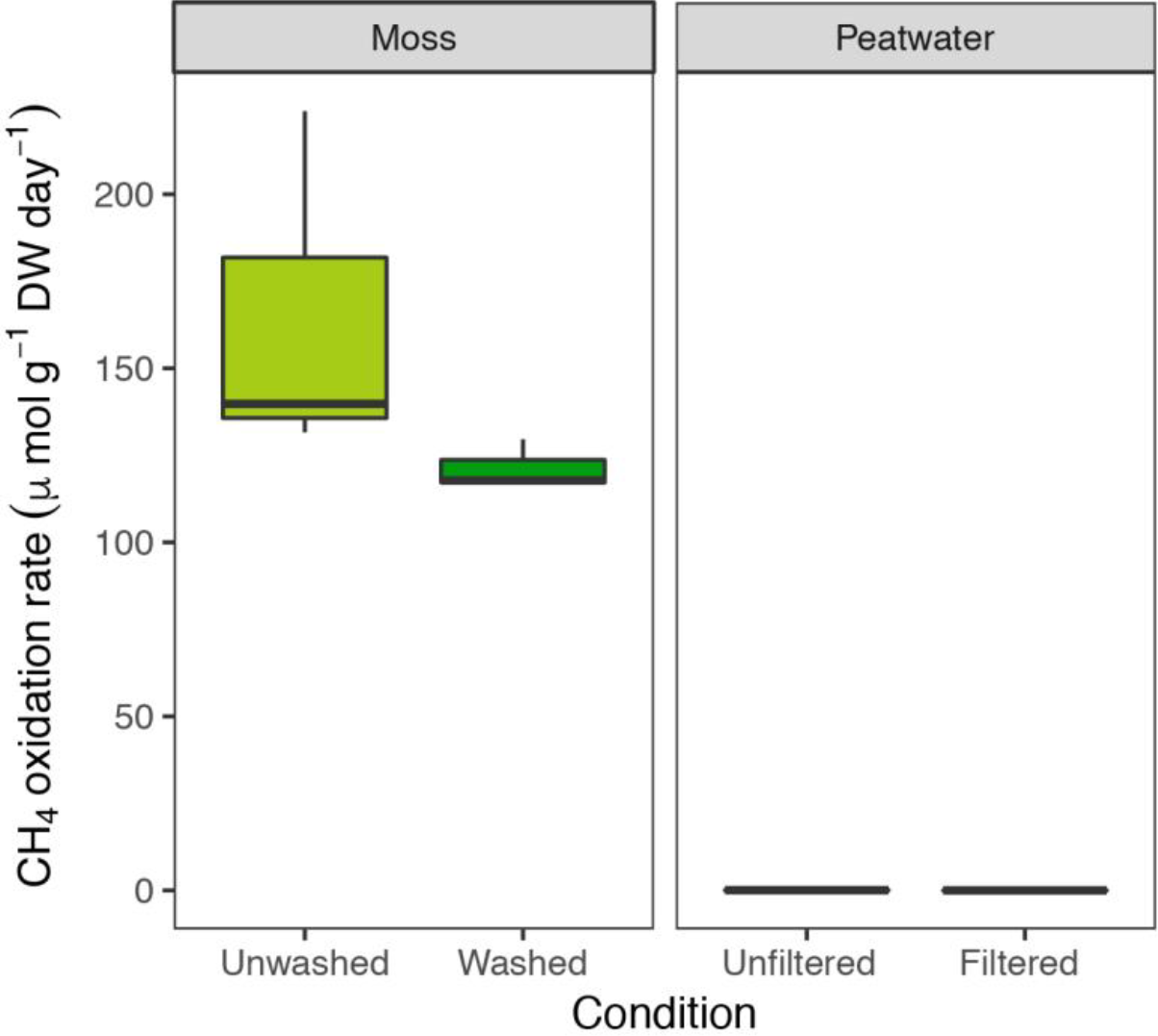
Potential CH_4_ oxidation rate in batch, associated with field *Sphagnum* mosses (light green, μmol CH_4_ g^−1^ DW day^−1^) or washed *Sphagnum* mosses (darker colors) and rates in peat water unfiltered or filtered. Error bars indicate the standard error of the mean (n=3).

### Mesocosm incubation

Two parallel mesocosm incubations were performed, one including a *Sphagnum* layer and one without. The net CH_4_ flux in the mesocosm showed a similar pattern for both mesocosms until day 8 of the incubation (Fig. 4). After 8 days, the moss mesocosm headspace always showed a lower CH_4_ concentration than the control mesocosm with only peat water. Furthermore, the emission from the *Sphagnum* moss mesocosm gradually decreased over the 32 day of the incubation, which is a strong indication of increasing CH_4_ oxidation activity. The variation in Fig. 4 is partly due to the daily manual refreshment of CH_4_ and air. The experiment was repeated for a second time, and the replicate incubation showed a similar pattern, with lower CH_4_ emission with the presence of *Sphagnum* moss layer (Supplementary Fig. S8 and Tables S7 and S8).

**Fig. 4.**
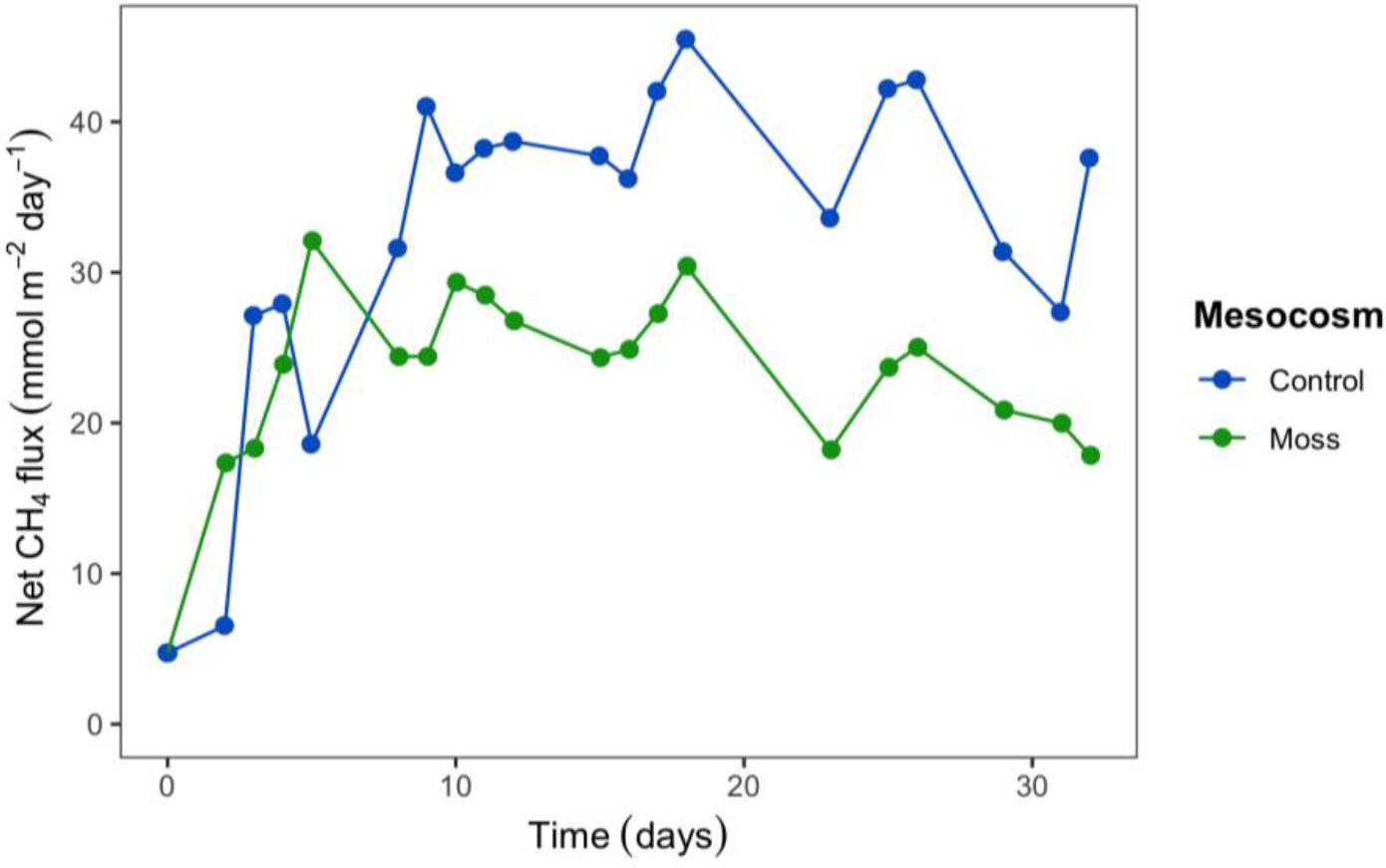
Net CH_4_ flux (mmol CH_4_ m^−2^ day^−1^) from the mesocosms with *Sphagnum* moss (green) and the control mesocosm with only peat water (blue) measured in the headspace over time (days). Each dot represents the mean of 2 technical replicates.

### Methane oxidation activity after mesocosm incubation

After 32 days of incubation in the mesocosms, the CH_4_ oxidation activity was determined in batch for each element of both mesocosms (water and/or moss). The CH_4_ oxidation activity was on average 189 μmol CH_4_ g^−1^ DW day^−1^ (Table 1) in mosses. Even after mesocosm incubation the peat water showed no CH_4_ oxidation activity (R^2^ <0.9; see Table 1 and Figs. S6 and S7), indicating that the water is not a favorable place for MOB. In the presence of acetylene, CH_4_ oxidation associated with the mosses was almost completely inhibited (F_(1,4)_ = 981.3, p < 0.001), indicating that the CH_4_ oxidation rate is entirely associated with methanotrophic microorganisms in or at the moss. Compared to the start of the incubation, the CH_4_ oxidation activity associated with mosses had increased by 155% (from 121 to 189 μmol g DW^−1^ day^−1^; Table 1 and Fig. 3).

**Table 1.** Potential CH_4_ oxidation rate in batch, after mesocosm incubation. Moss and peat water samples from each mesocosm were incubated in batch, with or without acetylene. Different italic letters indicate statistical differences between PMO rates, tested by 3-way Anova.

| Material | Mesocosm | Treatment | Potential methane oxidation rate<br>($\mu\text{mol CH}_4 \text{ g}^{-1} \text{ DW day}^{-1}$) | | SEM | R<sup>2</sup> | n |
| --- | --- | --- | --- | --- | --- | --- | --- |
| Moss | Moss |  | 189 | *a* | 6 | 0.98 | 3 |
| Moss | Moss | + acetylene | 2.0 | *b* | 2 | 0.30 | 3 |
| Material | Mesocosm | Treatment | Potential methane oxidation rate<br>($\mu\text{mol CH}_4 \text{ ml}^{-1} \text{ day}^{-1}$) | | SEM | R<sup>2</sup> | n |
| Water | Moss |  | 0.02 | *a* | 0.02 | 0.17 | 3 |
| Water | Moss | + acetylene | 0.03 | *a* | 0.01 | 0.51 | 3 |
| Water | Peatwater only |  | 0.09 | *b* | 0.01 | 0.86 | 3 |
| Water | Peatwater only | + acetylene | 0.05 | *b* | 0.01 | 0.67 | 3 |

### qPCR

To quantify the microbial community, both qPCR and amplicon sequencing of 16S rRNA genes were performed. Quantification of the bacteria (16S rRNA gene; Fig. 5) showed that bacterial copy numbers differed between all stages (F_(2,6)_=34.3, p<0.001). Substantial amounts (98%) of presumably epiphytes were washed away (Tukey HSD p<0.001). At the end of the incubations the copy numbers were back to about 97% of the original value (Tukey HSD p<0.05).

**Fig. 5.**
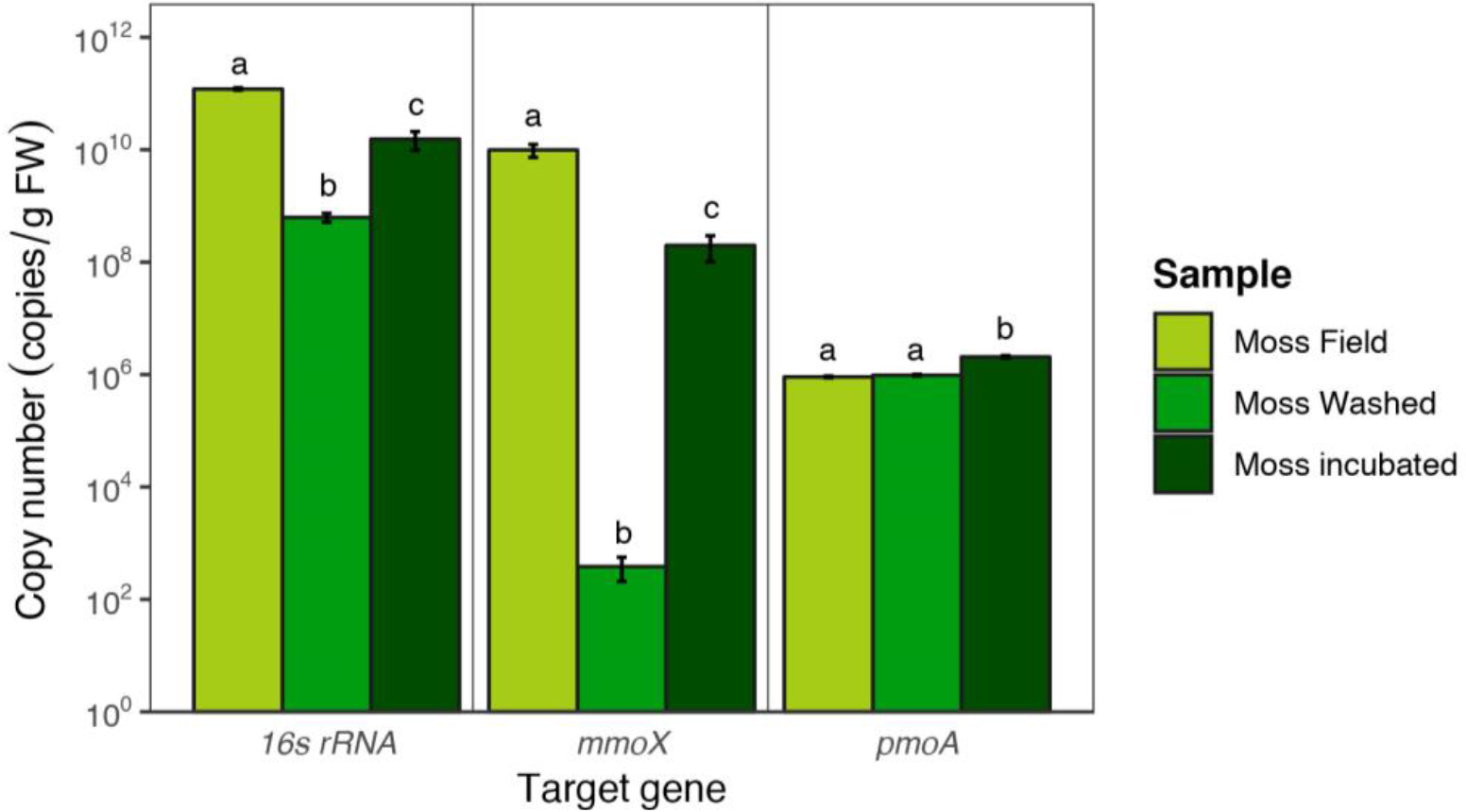
Copy numbers of bacteria 16S rRNA, *pmoA* and *mmoX* genes obtained via qPCR. Error bars indicate the standard error of the mean (n=3).

Quantification of methanotrophic microorganisms by *mmoX* gene and *pmoA* gene amplification showed a similar trend (*mmoX* F_(2,6)_=40.7, p<0.001; *pmoA* F_(2,6)_=27.1, p<0.001; Fig. 5). The *pmoA*-containing methanotrophs were overall less abundant than *mmoX*-containing methanotrophs (resp. 10^5^ vs. 10^10^ copies). The washing step greatly reduced the abundance of the *mmoX*-containing methanotrophs from 10^10^ to 10^2^ copies (Tukey HSD p<0.001), whereas *pmoA*-containing methanotrophs were much less affected (remained around 10^5^ copies; Tukey HSD p>0.05). Upon mesocosm incubation *mmoX* copies increased from 10^2^ to 10^8^ (Tukey HSD p<0.001), while *pmoA*-containing methanotrophs marginally increased from 10^5^ to 10^6^ copies (Tukey HSD p<0.01).

### Microbial community (16S rRNA gene)

The microbial community associated with the mosses was studied by 16S rRNA gene sequencing of the V3-V4 region. Comparison of the moss microbial community in the field, after washing and after incubation in the mesocosm shows a gradual change in microbial community. However, the main classes remained present throughout the incubation. Furthermore, mesocosm incubation increased diversity of the microbial community (Shannon and Chao 1 index, Table S4).

Looking at microbial community composition depicted as relative abundances in Fig. 6A, the *Proteobacteria* were the overall dominant phylum. Relative abundance of *Proteobacteria* was not affected by washing, but decreased during incubation in our mesocosm set-up. For the *Verrucomicrobiae* the relative abundance was lower after washing and increased after incubation. Especially the relative abundance of *Pedosphaerales* and *Opitutales* increased upon incubation (Supplementary Table S5). When focusing on the methanotrophic community, the relative abundance of Verrucomicrobial *Methylacidiphilales* increased by incubation (Fig. 6B). Other methanotrophic species whose relative abundance increased upon incubation are *Methylomonas* spp. and *Methylocystis* spp. (Fig. 6B). Only acidophilic *Methylocystis* isolates, *M. bryophila* and *M. heyeri* [53, 54], are known to contain both sMMO and pMMO, whereas neutrophilic *Methylocystis* and *Methylocella* species isolated so far only contain pMMO.

**Fig. 6.**
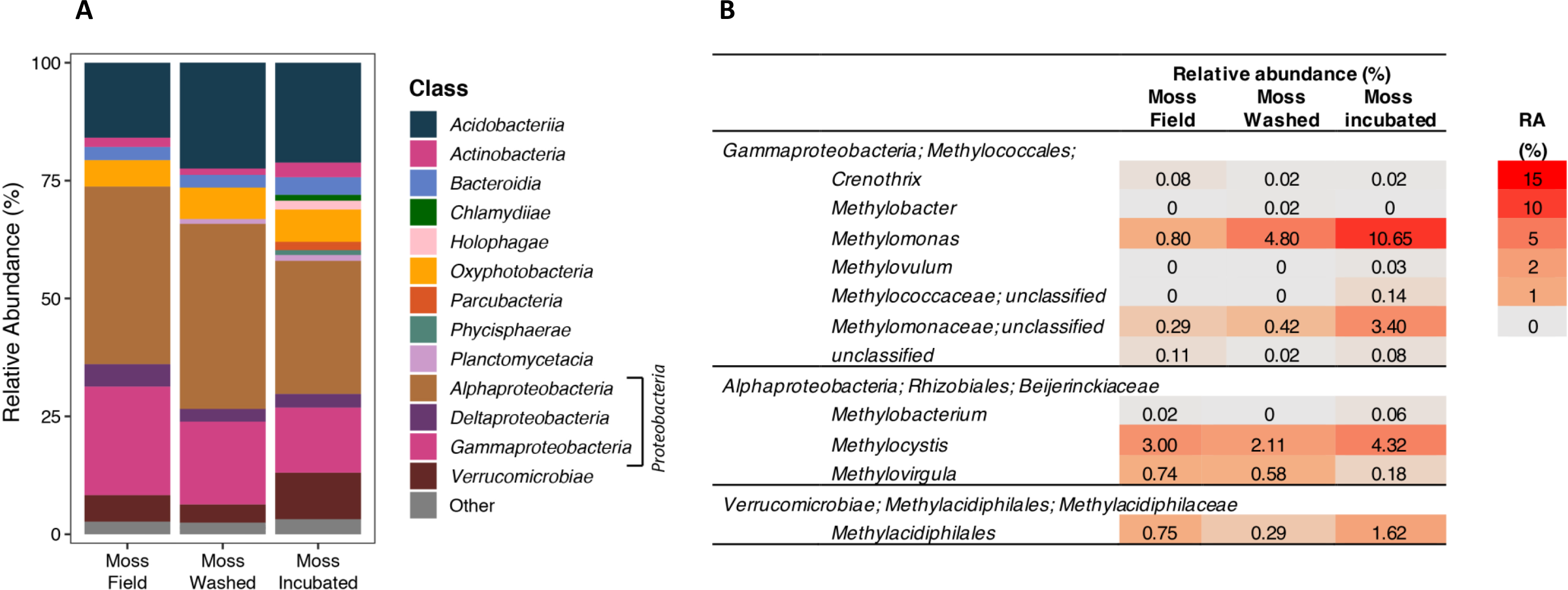
**A** Phylogenetic classification of the bacterial community based on 16S rRNA gene amplification and sequencing. Taxonomic groups with a relative abundance <1% are depicted as “Other”. In **B** specific relative abundances (RA in %) of methanotrophic bacteria in the bacterial 16s rRNA community profile are shown.

## Discussion

### Mesocosm approach

Studying and sampling the *Sphagnum* microbiome in the field is challenging, because the microbial community associated with the moss is influenced by many biotic and abiotic factors that are not controlled for. After many field campaigns we set out to circumvent these challenges and fluctuations. Therefore, we designed a novel mesocosm set up to mimic submerged *Sphagnum* moss ecosystem and operated it under controlled laboratory conditions to shed light on the association between aerobic CH_4_ oxidizers and a submerged *Sphagnum cuspidatum* community. We hypothesized that the submerged *Sphagnum* moss layer acts as a biofilter for CH_4_ and expected that the CH_4_-oxidizing community was mainly associated with *Sphagnum* moss. Indeed, in this controlled mesocosm set-up, we were able to mimic a significant reduction (31%) in CH_4_ emissions as was also observed in the field (Figs. 4 and S8). This CH_4_ removal was only associated with the mosses and not found in the peat water.

The novel mesocosm set-up allowed for enrichment of both methanotrophic activity and their abundance. Potential CH_4_ oxidation batch assays revealed a significant increase in methanotrophic activity after mesocosm incubation (from 121 ± 4 to 189 ± 6 μmol CH_4_ g^−1^ DW day^−1^, resp. Fig. 3 & Table 1). Similarly, qPCR of functional methanotrophic genes (*mmoX* and *pmoA*), indicated that significant numbers of CH_4_-oxidizing bacteria were present in and on the moss and that their numbers increased over the course of the incubation.

### Peat mosses strongly facilitate CH_4_-oxidizing activity

Washing of the moss and filtering of the peat water had little effect on CH_4_ oxidation activity and community composition, which underlines the tight association between CH_4_ oxidizers and *Sphagnum* mosses. Yet, qPCR revealed that bacterial copy numbers decreased by washing of the moss. The number of sMMO-containing methanotrophs decreased most significantly during washing, indicating that these methanotrophs might only be loosely attached epiphytes on the *Sphagnum* mosses. However, they showed the highest increase (10^2^ to 10^8^ copies/g FW) upon mesocosm incubation, equaling growth of up to 20 generations in 32 days. The transcription of *mmoX* gene and activity of sMMO-containing methanotrophs has previously been reported in peatlands [55–57]. The increase in sMMO copy number during incubation suggests that sMMO-containing methanotrophs are environmentally relevant in acidic peatland ecosystems, especially in submerged conditions, but their importance and contribution needs further study. Surprisingly, the pMMO-containing methanotrophs were initially less abundant than sMMO-containing methanotrophs, but seemed more tightly associated to the moss as washing had no effect on the copy numbers. There was hardly any increase in abundance upon incubation. Lack of copper might explain why pMMO containing methanotrophs did not thrive in the mesocosm incubation [20]. Ultimately, the enrichment of sMMO-containing methanotrophs upon mesocosm incubation shows that this set-up can be used to further study the functioning of sMMO methanotrophs in *Sphagnum* mosses as their ecology is far less understood than that of canonical pMMO containing methanotrophs.

### Microbial community composition

The *Sphagnum*-associated microbial community in all samples of this study showed high similarity to previous *Sphagnum*-associated 16S rRNA gene libraries [34, 38, 39]. Similar dominant community members were found in this study, with dominant phyla being the *Proteobacteria* (*Alpha*- and *Gammaproteobacteria*), *Cyanobacteria* (*Oxyphotobacteria*) and *Acidobacteria* and a relatively high abundance of *Verrucomicrobia*. Upon mesocosm incubation the microbial diversity increased, potentially due to the limited amount of nutrients present compared to field conditions. The relative abundance of *Verrucomicrobia* and *Planctomycetes* increased, whereas the relative abundance of the *Proteobacteria* decreased. Which processes control the changes in the moss-associated microbial community is topic for further study.

### Strong natural CH_4_ filter

The reduction of CH_4_ emission by the *Sphagnum*-methanotroph interaction in the studied mesocosm set-up is large (31%), compared to other high CH_4_ producing moss-dominated ecosystems. In other ecosystems CH_4_ oxidation also mitigates CH_4_ emission. For example, in the arctic tundra [28] 5% of the total CH_4_ emission is mitigated, whereas in hollows in *Sphagnum*-dominated peatland [58] measured CH_4_ production and oxidation rates and calculated that nearly 99% of the CH_4_ emission was mitigated by CH_4_-oxidizing microorganisms. For free-floating wetland plants, it was shown that up to 70% of the CH_4_ emission may be oxidized by the combination of decreased flux rates and high CH_4_-oxidizing activity [59].

Yet, the CH_4_ activity in the mesocosm set-up it is lower than the reduction found in the field. This is likely to be caused by the peat moss density, which was much higher in the field, where the moss layer was more than 50 cm deep. Although the stabilization of the net CH_4_ flux in both mesocosms occurred relatively quickly (8 days) and considerable CH_4_ mitigation was measured after 32 days of incubation, we believe that the CH_4_ mitigation by the moss associated methanotrophs in the mesocosm will most probably increase even further by prolonging the incubation time and increased amount of *Sphagnum* mosses. Additionally, the mesocosm set-up could be improved by replacing the manual addition of CH_4_ and air of the mesocosm with a continuous supply system. In a continuous bioreactor set-up, the system is even more stable, and variation is further reduced. The high reduction in CH_4_ emission in submerged *Sphagnum* emphasizes that the methanotrophs associated with *Sphagnum* are important in CH_4_ cycling in peatlands [12, 28–30], as they strongly regulate CH_4_ emission from *Sphagnum* dominated peatlands.

### Implications for degraded peatlands

The large organic matter stocks in peatlands are a great potential source for CO_2_ when these peatlands are drained. Restoration measures aimed at preventing further oxidation and degradation of these drained peatlands, often involve hydrological measures (rewetting), resulting in inundation of large surface areas. After rewetting, anaerobic degradation of organic matter will result in high CH_4_ production rates. As shown above, methanotrophs are tightly associated to *Sphagnum* mosses. Presence of this consortium in restored peatlands can thereby strongly mitigate CH_4_ emissions. Since the presence and abundance of *Sphagnum* in peatlands is affected by peatland degradation as well [60, 61], care should be taken to restore and facilitate *Sphagnum* mosses in restored peatlands.

### Conclusion

*Sphagnum* mosses have many key roles in peat ecosystems [62], and this study shows that their microbiome and specifically the methanotrophs associated with *Sphagnum* are crucial to keep CH_4_ emissions from *Sphagnum*-dominated peatlands low. Peatland restoration practices involving rewetting, should therefore aim to stimulate *Sphagnum* growth simultaneously, in order to keep CH_4_ emissions at bay. The presented mesocosm set-up can be used to further study the effect of various climate change relevant factors (such as temperature, pH, fertilization) on CH_4_ cycling in submerged *Sphagnum* moss ecosystems. Studying the influence of climate change on the *Sphagnum*-methanotroph interaction and CH_4_ balance is crucial to get a better understanding of the potential positive feedback loop that reside in peatlands.

## Acknowledgements

We thank Nardy Kip for support with the initial design of the mesocosms. Tijs van den Bosch is thanked for helping out with the 16S rRNA sequencing. The General Instruments department at the Faculty of Science at Radboud University, especially Paul van der Ven and Sebastian Krosse are thanked for measuring elemental composition of the water samples and determining stable isotope contents.

## Availability of data and materials

All sequencing data has been deposited in the NCBI SRA database, project number PRJNA517391.

