## Supplementary tables and figures for "A novel mesocosm set-up reveals strong methane emission reduction in submerged peat moss *Sphagnum cuspidatum* by tightly associated methanotrophs"

### Supplementary material

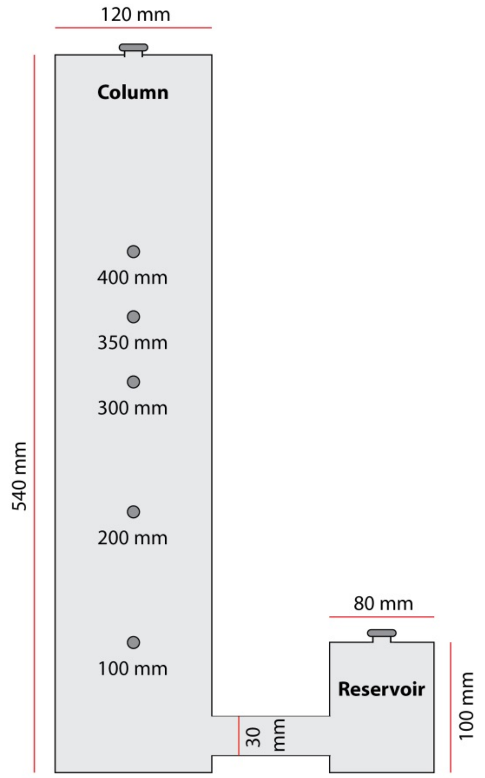

**Fig. S1** Dimensions of mesocosm columns.

**Table S1** Change in read count during data processing.

| Step | | Moss Field | | | Moss Washed | | | Moss incubated | | |
|---|---|---|---|---|---|---|---|---|---|---|
| | | Read count (average) | | SEM | Read count (average) | | SEM | Read count (average) | | SEM |
| 1 | Unprocessed reads | 31 301 | ± | 2 569 | 32 001 | ± | 2 117 | 45 380 | ± | 13 738 |
| 2 | Screen sequences | 28 443 | ± | 2 351 | 29 082 | ± | 1 958 | 41 585 | ± | 12 791 |
| 3 | Unique sequences | 27 912 | ± | 2 311 | 28 488 | ± | 1 923 | 40 541 | ± | 12 439 |
| 4 | Aligned sequences | 27 912 | ± | 2 311 | 28 488 | ± | 1 923 | 40 541 | ± | 12 439 |
| 5 | Screened alignment | 27 912 | ± | 2 311 | 28 488 | ± | 1 923 | 40 541 | ± | 12 439 |
| 6 | Filter positions in alignment | 27 912 | ± | 2 311 | 28 488 | ± | 1 923 | 40 541 | ± | 12 439 |
| 7 | Pre-clustering | 27 912 | ± | 2 311 | 28 488 | ± | 1 923 | 40 541 | ± | 12 439 |
| 8 | Chimera removal | 23 924 | ± | 2 032 | 24 129 | ± | 1 667 | 31 278 | ± | 9 097 |
| 9 | Non-target lineage removal | 7 847 | ± | 578 | 9 028 | ± | 669 | 18 003 | ± | 5 700 |
| | *Total read count per group* | *23 542* | | | *27 083* | | | *54 009* | | |
| 10 | Random subsample | 6 500 | ± | 0 | 6 500 | ± | 0 | 6 500 | ± | 0 |

**Table S2** Change in read quality during data processing.

| Step | | Total read count | # Unique reads | Mean read length (bp) | Max. ambiguent positions | Max. polymers |
|---|---|---|---|---|---|---|
| 1 | Unprocessed reads | 326 045 | | 444 | 308 | 297 |
| 2 | Screen sequences | 297 333 | | 445 | 0 | 45 |
| 3 | Unique sequences | 297 333 | 154 224 | 445 | 0 | 45 |
| 4 | Align sequences | 297 333 | 154 224 | 407 | 0 | 28 |
| 5 | Screen alignment | 290 823 | 148 011 | 407 | 0 | 8 |
| 6 | Filter positions in alignment | 290 823 | 82 822 | 407 | 0 | 8 |
| 7 | Pre-clustering | 290 823 | 50 548 | 407 | 0 | 8 |
| 8 | Chimera removal | 237 994 | 12 021 | 407 | 0 | 8 |
| 9 | Non-target lineage removal | 104 634 | 10 068 | 411 | 0 | 8 |

**Table S3** Primer sequences used in this study.

| Target gene | Primer name | Sequence | Reference |
|---|---|---|---|
| *pmoA* | A189f | 5'-GGNGACTGGGACTTCTGG-3' | (Holmes et al. 1995) |
| | A682r | 5'-GAASGCNGAGAAGAASGC-3' | (Holmes et al. 1995) |
| *mmoX* | mmoX1 | 5'-CGGTCCGCTGTGGAAGGGCATGAAGCGCGT-3' | (Miquez et al. 1997) |
| | mmoX2 | 5'-GGCTCGACCTTGAACTTGGAGCCATACTCG-3' | (Miquez et al. 1997) |
| *16S rRNA* | Bact 341f | 5′-CCTACGGGNGGCWGCAG-3′ | (Klindworth et al. 2013) |
| *Bacteria* | Bact 785r | 5'-GACTACHVGGGTATCTAATCC-3 | (Klindworth et al. 2013) |

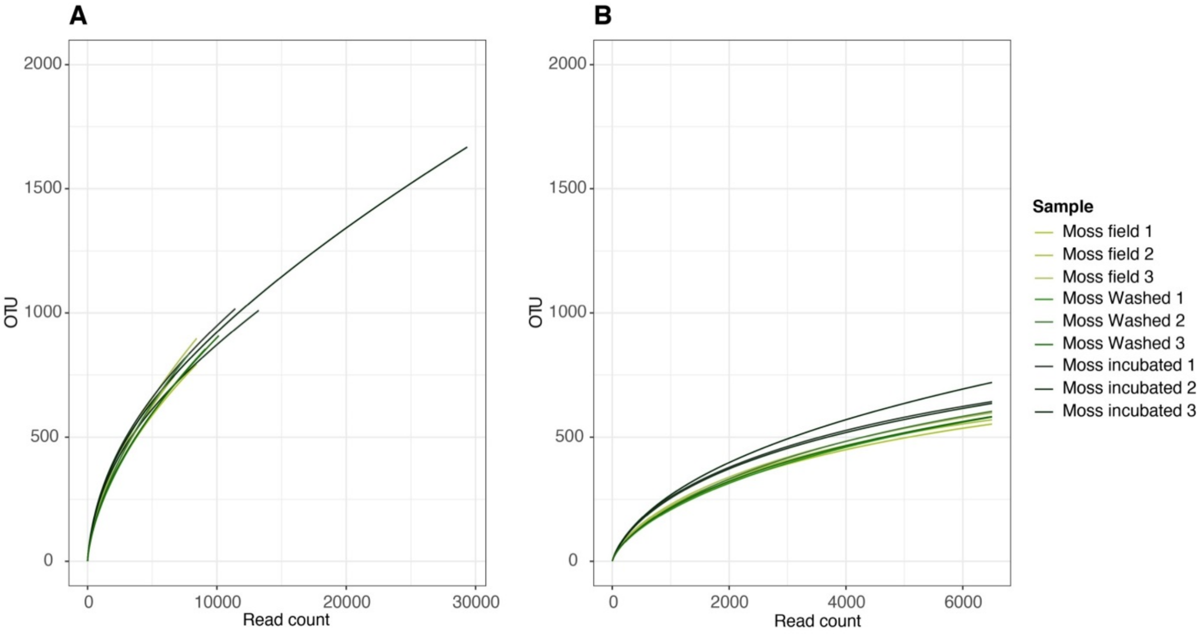

**Fig. S2** Rarefaction curves before **(A)** and after **(B)** subsampling, singletons were removed.

**Table S4** Alpha diversity index for the different samples.

| | | Richness | | | | Evenness |
|---|---|---|---|---|---|---|
| **Sample** | **Observed species number (S)** | **Shannon (H)** | **Fisher's alpha** | **Chao1** | **ACE** | **Pielou (J)** |
| **Moss Field** | 1136 | 4.53 | 263.0 | 1607.3 | 1729.4 | 0.64 |
| **Moss Washed** | 1215 | 4.23 | 287.0 | 1762.1 | 2001.5 | 0.60 |
| **Moss Incubated** | 1213 | 4.77 | 286.4 | 2046.3 | 2061.7 | 0.67 |

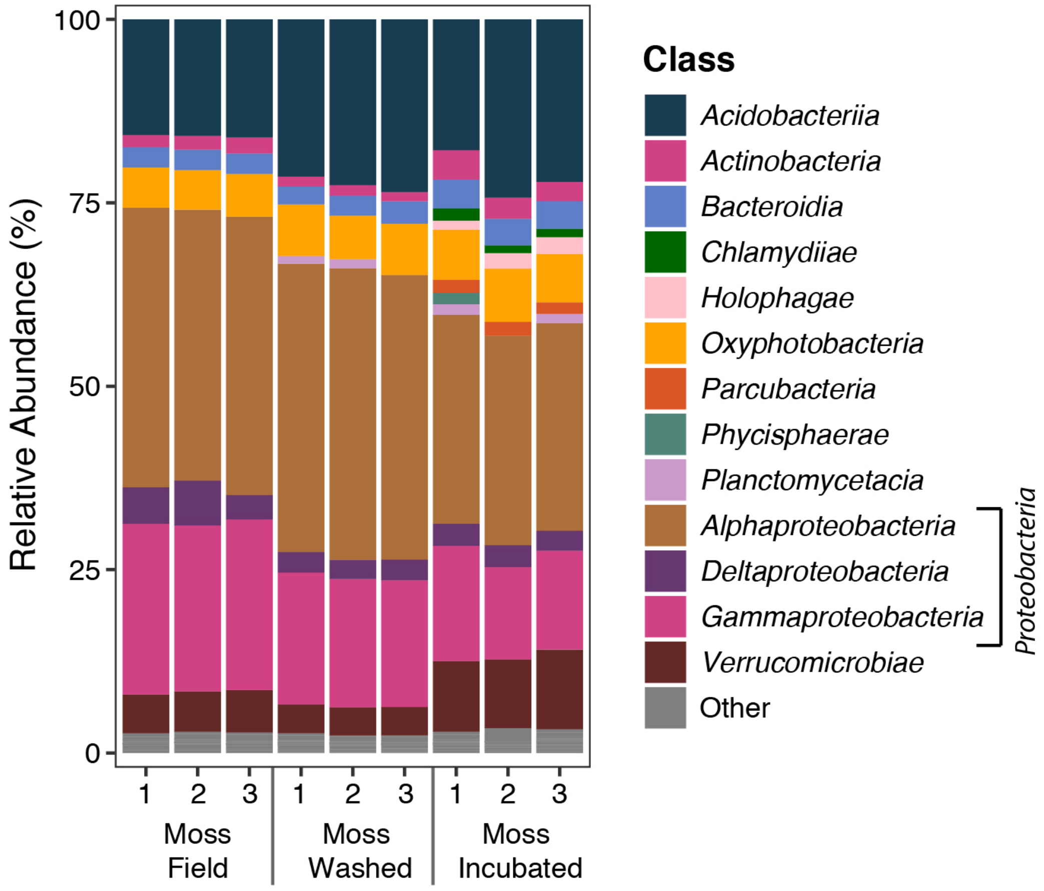

**Fig. S3** Phylogenetic classification of the bacterial community per sample based on 16s rRNA gene amplification and sequencing. Taxonomic groups with a relative abundance <1% are depicted as "Other".

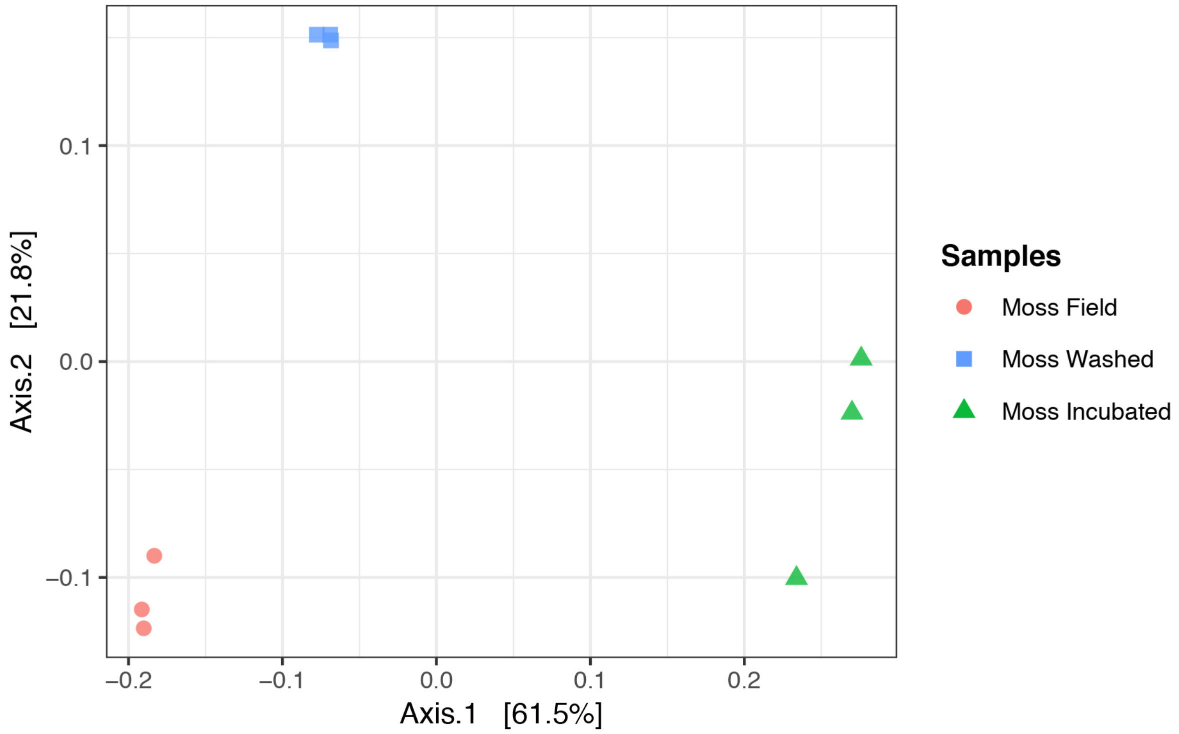

**Fig. S4** Principal Coordinate Analysis (PCoA) of the microbial diversity using Bray Curtis dissimilarity. The different samples are coded according to colour.

**Tables S5** Relative abundances (RA in %) of Verrucomicrobial microorganisms in the bacterial 16s rRNA community profile.

| | **Relative abundance (%)** | | |
|---|---|---|---|
| | **Moss Field** | **Moss Washed** | **Moss incubated** |
| *Verrucomicrobia; Verrucomicrobiae;* | | | |
| *Chthoniobacterales* | 6.60 | 3.11 | 3.15 |
| *Methylacidiphilales* | 0.75 | 0.29 | 1.62 |
| *Opitutales* | 2.06 | 2.78 | 7.75 |
| *Pedosphaerales* | 6.54 | 4.23 | 14.85 |
| *S-BQ2-57_soil_group* | 0.02 | 0 | 0 |
| *unclassified* | 0.02 | 0.15 | 0.08 |
| *Verrucomicrobiales* | 0.09 | 0.06 | 0.94 |

| **RA (%)** |
|---|
| 15 |
| 10 |
| 5 |
| 2 |
| 1 |
| 0 |

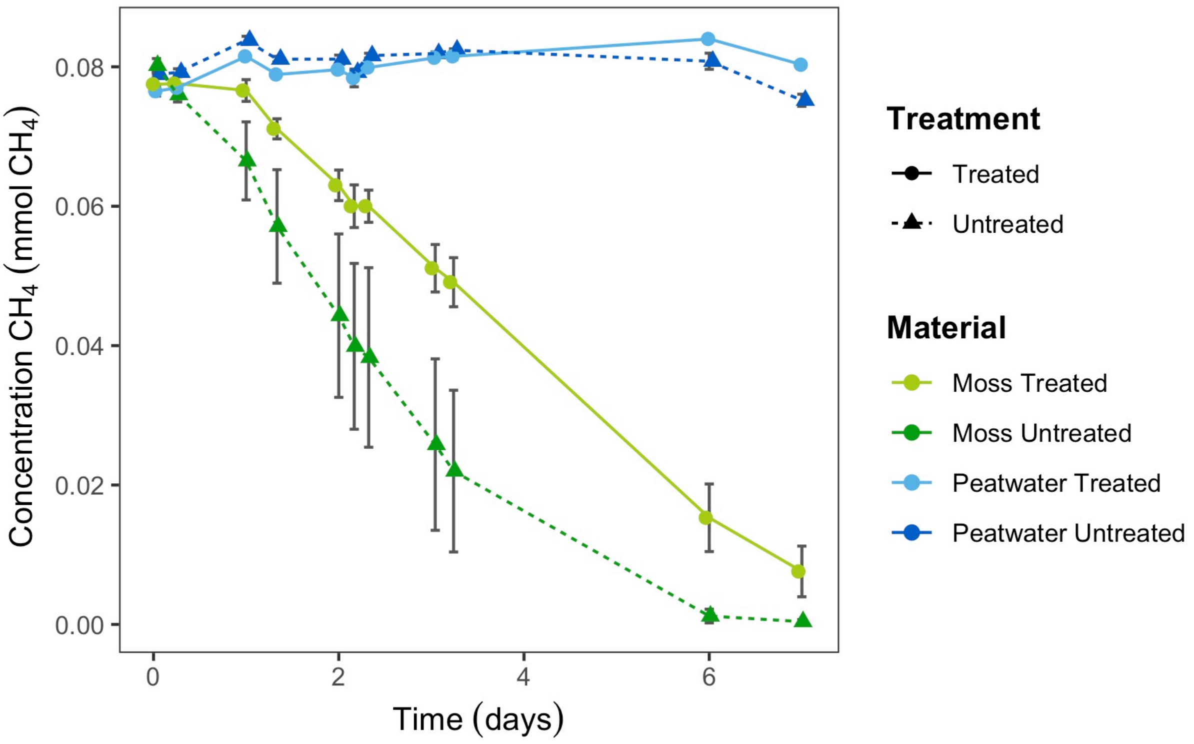

**Fig. S5** Potential $CH_4$ oxidation rate in batch assays, prior to mesocosm incubation round 1. $CH_4$ concentration (mmol) in the headspace is displayed over time (days) for incubations with: mosses (unwashed (dark green, n=3) and washed (light green, n=3)) and peat water (unfiltered (dark blue, n=3) and filtered (light blue, n=3)). Error bars indicate the standard error of the mean.

**Table S6** Mean potential $CH_4$ oxidation rate in mosses (µmol $CH_4$ $g^{-1}$ DW $day^{-1}$) and peat water (µmol $CH_4$ $ml^{-1}$ $day^{-1}$) prior to mesocosm incubation round 1.

| ***Round 1*** | | | | | |
|---|---|---|---|---|---|
| **Material** | **Treatment** | **Methane oxidation rate** (µmol $CH_4$ $g^{-1}$ DW $day^{-1}$) | **SE** | **R2** | **n** |
| **Moss** | Washed | 121 | 4.1 | 0.98 | 3 |
| **Moss** | Field | 165 | 29 | 0.99 | 3 |
| **Material** | **Treatment** | **Methane oxidation rate** (µmol $CH_4$ $ml^{-1}$ $day^{-1}$) | **SE** | **R2** | **n** |
| **Water** | Filtered | 0.01 | 0.003 | 0.40 | 3 |
| **Water** | Field | 0.08 | 0.02 | 0.48 | 3 |

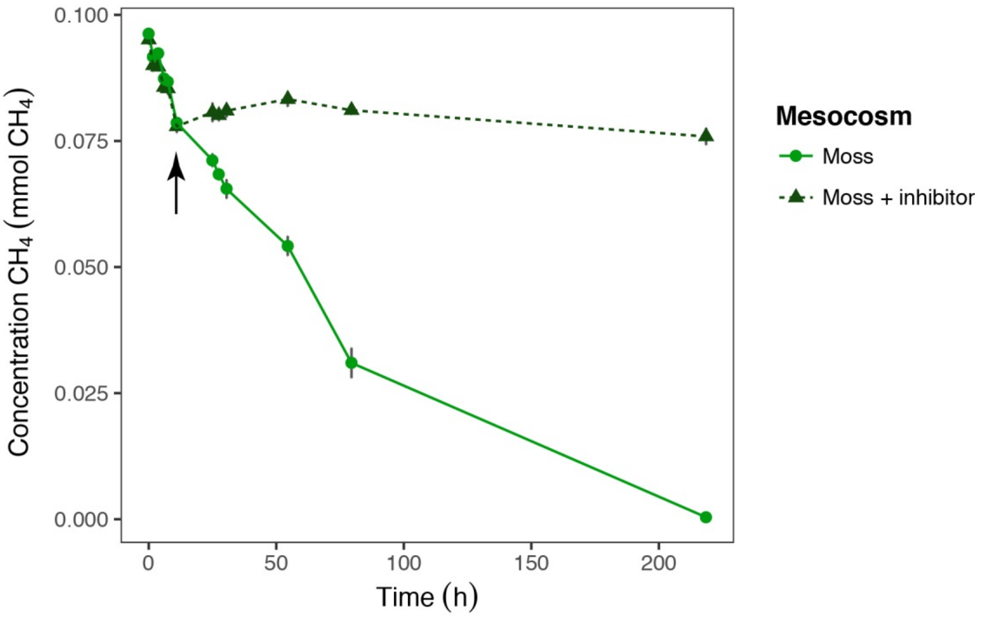

**Fig. S6** Potential $CH_4$ oxidation rate of the mosses in batch assays, after mesocosm incubation round 1. $CH_4$ concentration (mmol) in the headspace is displayed over time (h) for incubations with mosses, either with PMO inhibitor acetylene (light green, n=3) or without PMO inhibitor (dark green, n=3). Arrow indicates addition of acetylene gas. Error bars indicate the standard error of the mean.

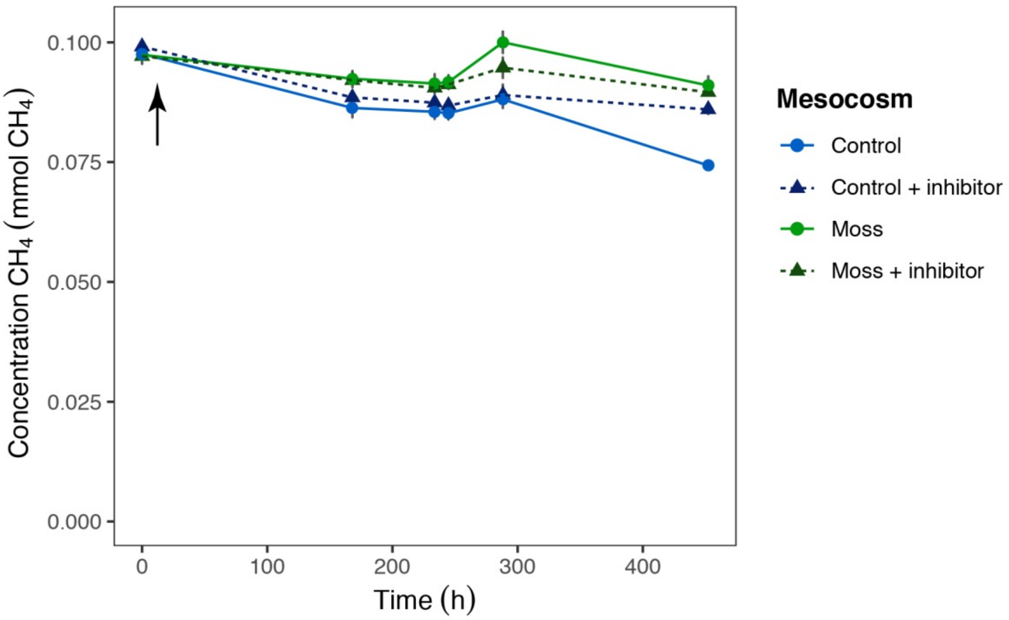

**Fig. S7** Potential $CH_4$ oxidation rate of peat water in batch assays, after mesocosm incubation round 1. $CH_4$ concentration (mmol) in the headspace is displayed over time (h) for incubations with peat water from the moss mesocosm (green) or peat water from the control mesocosm (blue). Incubations are either with PMO inhibitor (lighter colors, dashed line, n=3) or without PMO inhibitor (darker colors, dashed line, n=3). Arrow indicates addition of acetylene gas. Error bars indicate the standard error of the mean.

**Table S7** Mean potential $CH_4$ oxidation rate in mosses (µmol $CH_4$ $g^{-1}$ DW $day^{-1}$) and peat water (µmol $CH_4$ $ml^{-1}$ $day^{-1}$) prior to mesocosm incubation round 2.

| ***Round 2*** | | | | | |
|---|---|---|---|---|---|
| **Material** | **Treatment** | **Methane oxidation rate** (µmol $CH_4$ $g^{-1}$ DW $day^{-1}$) | **SE** | **R2** | **n** |
| **Moss** | Washed | 215 | 7.6 | 0.99 | 3 |
| **Moss** | Field | 281 | 14 | 0.99 | 3 |
| **Material** | **Treatment** | **Methane oxidation rate** (µmol $CH_4$ $ml^{-1}$ $day^{-1}$) | **SE** | **R2** | **n** |
| **Water** | Filtered | -0.04 | 0.03 | 0.35 | 3 |

**Table S8** Mean potential $CH_4$ oxidation rate in mosses (µmol $CH_4$ $g^{-1}$ DW $day^{-1}$) and peat water (µmol $CH_4$ $ml^{-1}$ $day^{-1}$) after mesocosm incubation round 2.

| ***Round 2*** | | | | | | |
|---|---|---|---|---|---|---|
| **Material** | **Mesocosm** | **Treatment** | **Methane oxidation rate** (µmol $CH_4$ $g^{-1}$ DW $day^{-1}$) | **SE** | **R2** | **n** |
| **Moss** | Moss | | 121 | 12 | 0.97 | 3 |
| **Moss** | Moss | + inhibitor | -2.0 | 0.6 | 0.33 | 3 |
| **Material** | **Mesocosm** | **Treatment** | **Methane oxidation rate** (µmol $CH_4$ $ml^{-1}$ $day^{-1}$) | **SE** | **R2** | **n** |
| **Water** | Moss | | 0.10 | 0.02 | 0.94 | 3 |
| **Water** | Moss | + inhibitor | -0.02 | 0.00 | 0.71 | 3 |
| **Water** | Control | | -0.01 | 0.01 | 0.39 | 3 |
| **Water** | Control | + inhibitor | -0.02 | 0.00 | 0.91 | 3 |

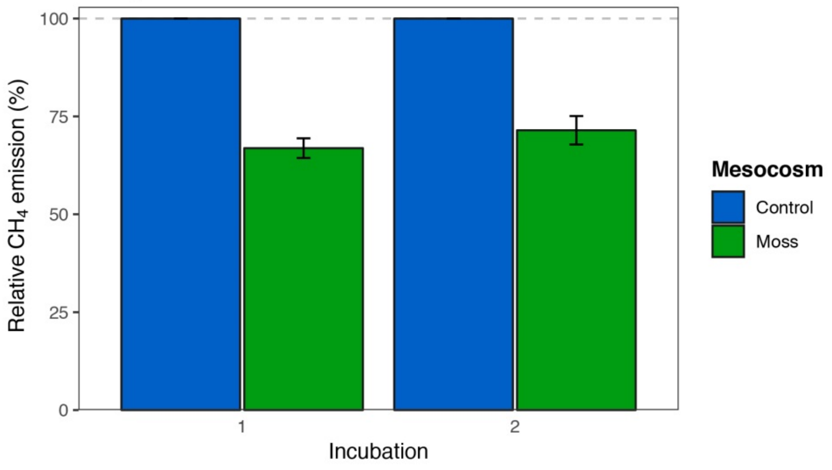

**Fig. S8** Relative $CH_4$ emission (as % of emission in control mesocosm) from the control mesocosm (blue) and the moss mesocosms (green), shown for incubation round 1 and incubation round 2. Each bar represents the average relative $CH_4$ emission of days 8-32 and error bars indicate the standard error.

**Table S9** Composition of the unfiltered and filtered porewater.

| porewater | $NO_3^-$ (µmol $L^{-1}$) | | | $NH_4^+$ (µmol $L^{-1}$) | | | $PO_4^{3-}$ (µmol $L^{-1}$) | | | Al (µmol $L^{-1}$) | | | Ca (µmol $L^{-1}$) | | | Fe (µmol $L^{-1}$) | | | K (µmol $L^{-1}$) | |
|---|---|---|---|---|---|---|---|---|---|---|---|---|---|---|---|---|---|---|---|---|
| unfiltered | 1.22 | ± | 0.07 | 12.58 | ± | 2.83 | 0.68 | ± | 0.05 | 9.27 | ± | 2.21 | 30.51 | ± | 1.06 | 14.69 | ± | 2.45 | 9.03 | ± |
| filtered | 2.17 | ± | 0.54 | 3.55 | ± | 0.27 | 0.31 | ± | 0.06 | 4.52 | ± | 0.36 | 33.25 | ± | 2.06 | 4.12 | ± | 0.17 | 5.18 | ± |
| **porewater** | **Mg** (µmol $L^{-1}$) | | | **Mn** (µmol $L^{-1}$) | | | **Na** (µmol $L^{-1}$) | | | **P** (µmol $L^{-1}$) | | | **S** (µmol $L^{-1}$) | | | **Si** (µmol $L^{-1}$) | | | **Zn** (µmol $L^{-1}$) | |
| filtered | 26.62 | ± | 0.67 | 0.44 | ± | 0.02 | 172.44 | ± | 3.32 | 2.04 | ± | 0.21 | 24.08 | ± | 0.58 | 9.45 | ± | 0.40 | 0.58 | ± |
| unfiltered | 21.70 | ± | 0.04 | 0.46 | ± | 0.02 | 170.04 | ± | 3.28 | 0.48 | ± | 0.12 | 20.61 | ± | 0.67 | 9.85 | ± | 0.21 | 0.43 | ± |
